# Modeling alcoholic liver disease in a human Liver-Chip

**DOI:** 10.1101/2020.07.14.203166

**Authors:** Janna C. Nawroth, Debora B. Petropolis, Dimitris V. Manatakis, Tengku Ibrahim Maulana, Gabriel Burchett, Katharina Schlünder, Anke Witt, Abhishek Shukla, Geraldine Hamilton, Ekihiro Seki, Shelley Lu, Katia Karalis

## Abstract

Fatty liver disease (FLD), is a major public health burden that affects up to 30% of people in Western countries and leads to progressive liver injury, comorbidities, and increased mortality. Key risk factors for developing FLD are obesity and alcohol consumption, both of which are growing in prevalence worldwide. There is an urgent need for human-relevant preclinical models to improve our understanding of FLD progression to steatohepatitis and for the development of sensitive noninvasive diagnostics and therapies. Alcohol-induced liver disease (ALD) represents an ideal case for modeling FDL as ethanol exposure is a comparatively simpler trigger for experimental induction of the pathology, as opposed to the complexity of modeling the diet- and life-style induced FLD. Further, despite their different root causes, the two pathologies share several common characteristics in disease progression and deterioration of liver function, highlighting the potential of an ALD microphysiological model for broad application in translational research. Here, we leverage our recently reported human Liver-Chip for toxicity applications, to expand the capabilities of the platform for broad application in translational research. We report the first *in vitro* modeling of ALD that uses human relevant blood alcohol concentrations (BAC) and affords multimodal profiling of clinically relevant endpoints. Our ALD Liver-Chip recapitulates established FLD markers in response to ethanol in a concentration-dependent manner, including lipid accumulation and oxidative stress. Importantly, we show that the ALD Liver-Chip supports the study of secondary insults common in patients with advanced ALD, such as high blood endotoxin levels due to alcohol-associated increased intestinal permeability and barrier dysfunction. Moreover, owing to new developments in the design, the ALD Liver-Chip enables the measurement of structural changes of the bile canaliculi (BC) network as a novel *in vitro* quantitative readout of alcoholic liver toxicity. In summary, we report the development of a human ALD Liver-Chip as a new platform for modeling the progression of alcohol-induced liver injury with direct translation to clinical research.

## Introduction

Fatty liver disease (FLD) is a growing global health problem that affects up to 30% of the general population in Western countries^1–3^. The disease can be classified based on the causative trigger as either alcohol-associated liver disease (ALD) induced by excessive alcohol consumption, or as diet-induced non-alcoholic fatty liver disease/nonalcoholic steatohepatitis (NAFLD/NASH)^2^. Despite their differences in etiology and epidemiology, both forms of FLD share several genetic susceptibility markers, including PNPLA3, TM6SF2, MBOAT7 ^4,5^, and both progress from simple hepatic steatosis to liver fibrosis, cirrhosis, and finally hepatocellular carcinoma^6–11^. As reported recently, the worldwide prevalence of NAFLD and ALD was 25% and 4.6%, respectively^12^. More importantly, in the Western countries, including the US, ALD accounts for 45-50% of cirrhosis-associated death^13^. Excessive use of alcohol, a major social problem^14^, contributes substantially to the global burden of FLD. Specifically, of the 1% of the deaths due to liver cirrhosis, the end stage of FLD, ca. 50% are directly due to alcohol abuse^8,9,15^.

All the above highlight the urgency for the development of specific therapies and identification of noninvasive biomarkers of disease progression. For ALD, alcohol intake is the established cause of clinical ASH. Thus, according to the American Gastroenterological Association and the European Association for the Study of the Liver, in patients presenting with FLD, a history of alcohol intake above 20g/d for women and 30g/d for men suffices for making the diagnosis of ALD/ASH^2^. However, not all heavy alcohol drinkers cause alcohol-associated liver injury and steatohepatitis. Therefore, it is better to understand the underlying mechanism of the progression of alcohol-induced fatty liver to alcohol-associated steatohepatitis. Here, we hypothesized that modeling ALD may represent an ideal proof-of-concept study to test the capacity of the Liver-Chip to recapitulate development and progression from alcohol-induced fatty liver to alcohol-associated steatohepatitis. Notably, no human *in vitro* models for alcohol-induced steatosis using clinically relevant blood alcohol concentrations (BAC) exist.

The great majority of the experimental studies on FLD have employed animal models that display aspects of the liver disease phenotype but do not capture the spectrum of the metabolic, inflammatory and fibrotic responses found in the human patients^16,17^. Similarly, *in vitro* models such as hepatocyte sandwich culture and liver spheroids may recapitulate several of the features of the liver diseases but are missing the dynamics of the tissue microenvironment and the associated cell-cell interactions. A relevant example is given by the ethanol-induced alterations in hepatic cytochrome P450 CYP2E1 and their role in the mechanisms driving liver damage^18^. This mechanism so far can only by studied in human microphysiological systems as the expression of these enzymes is not maintained in commonly available liver cell lines (e.g., HepG2, HuH7 cells) as well as standard sandwich liver cell culture beyond the early time of the culture^19^. Overuse of alcohol in humans induces systemic effects, such as compromised intestinal epithelial barrier function and increased gut permeability (“leaky gut”), a response subject to significant species-dependent variability^20^. Leaky gut and the ensuing inflammatory response due to the escape of bacterial endotoxins to the portal and systemic circulation are suggested to act as the second factor, together with FLD, in the “two-hit” hypothesis for the development of ASH^20^.

All the above indicate the urgency for *in vitro* platforms that can more closely approach human organ physiology to enable basic research and translation to the clinic and eliminate the comorbidities of alcoholic patients, including a staggering number of deaths due to liver cirrhosis^21^. To address the need for in vitro platforms of human liver diseases, we developed a microphysiological system to enable the modeling of the pathogenesis and progression of ALD, based on clinically relevant endpoints. We have recently reported the development of a Liver-Chip that uses primary human cells to recreate the liver sinusoid architecture and we demonstrated its unique usefulness for the assessment of drug safety and toxicity for humans^19^. Here, we leveraged this Liver-Chip model and advanced the design with new capabilities to recapitulate critical events in human ALD using alcohol at concentrations as those found in the blood (BAC) of patients with ALD. Further, we evaluated the ability of the ALD Liver-Chip to model disease recovery through alcohol abstinence, as well as the worsening of the phenotype in a two-hit approach by co-exposure to alcohol and the Gram-negative bacterial endotoxin, lipopolysaccharide (LPS). We report here our findings based on multimodal profiling of the histological, molecular, and functional tissue changes in response to ethanol alone, or in combination with LPS, or following abstinence from ethanol. Our findings demonstrate the induction of ALD/ASH phenotypes supported by several metabolic stress-associated endpoints in the ALD Liver-Chip. In addition, we report disease-induced changes in the structure of the bile canaliculi (BC) network, representing a new readout of liver injury which has not been shown *in vitro* before although BC network structure has been reported as a sensitive marker of toxicity in mice^22^ and was also altered in human NAFLD patients^23^. For this purpose, we optimized the extracellular matrix (ECM) scaffolding to support robust formation of a biomimetic, highly branched BC network, and we captured the ethanol-induced disruptions of the BC network through novel quantitative imaging tools. Together, our results demonstrate the unique ability of the ALD Liver-Chip to model human relevant alcoholic steatosis and phenotypes previously missed by *in vitro* systems. We believe that the ALD Liver-Chip is a powerful platform for probing the mechanisms of steatosis onset and progression to steatohepatitis, and for supporting drug discovery by providing efficacy and safety endpoints in a patient-specific manner.

## Results

### Development of the Liver-Chip for modeling ALD/ASH

Our approach f’1234ertor modelling human ALD was to continuously perfuse our recently developed Liver-Chip, which is an organotypic microphysiological system (MPS) using primary human cells^19^, with ethanol doses within clinically relevant blood alcohol concentrations (BACs) (Fig. 1a), followed by multimodal phenotyping and functional analysis. As reported^19^, the Liver-Chip is made of polydimethylsiloxane (PDMS) and contains an upper channel (1 mm tall × 1 mm wide) and a lower channel (0.2 mm tall × 1 mm wide). The channels are separated by an ECM-coated PDMS membrane with 7-µm pores to allow for cell-cell interactions. The Tri-culture configuration used in this study includes primary hepatocytes cultured in the upper channel and primary liver sinusoidal endothelial cells (LSECs) cultured on the opposite side of the membrane, i.e. in the lower channel, along with Kupffer cells. As this design recapitulates key elements of the hepatic sinusoid microenvironment^19^, we were also particularly interested in recreating the bile canaliculi (BC) network that is established between neighboring hepatocytes. In mice, changes to the BC network structure and associated biliary clearance function have been shown to be an early and sensitive indicator of drug toxicity in the liver microenvironment^22^. However, in contrast to rodent hepatocytes, human hepatocytes do not readily form biomimetic bile canaliculi networks *in vitro*^24–28^. The latter together with the lack of a reproducible, quantitative manner to assess the integrity of the BC network have precluded the use of this clinical endpoint in preclinical studies. Notably, BC integrity is strongly dependent on the presence of a stable extracellular matrix that affords the symmetric mechanical anchoring needed for hepatocyte polarization and BC lumen formation^29^. Based on these findings, we assessed the impact of various ECM scaffold compositions and patterning methods on BC integrity by using quantitative metrics to describe the MRP-2 stained BC network topology (Fig. 1b and c and Supplementary Fig. 1A-C). In all the experiments for this study we used the matrix protocols “ECM-D” or “ECM-E” (see Methods for details) which promoted biomimetic BC network formation and resulted in the best and reproducible BC metrics (Fig 1b). In these ECM conditions, the primary hepatocytes were covered in a 150-300 µm thick 3D gel of Col-I^30^ (Supplementary Fig. 1C). Thus, by optimizing the Liver-Chip ECM scaffolding, we significantly improved formation of robust bile canaliculi networks in human hepatocytes. Further, we developed a digital pathology method for the quantitative assessment of BC integrity using structural metrics of the network geometry (see Methods), a sensitive method for the characterization of the Liver-Chip responses to ethanol.

**Figure 1:**
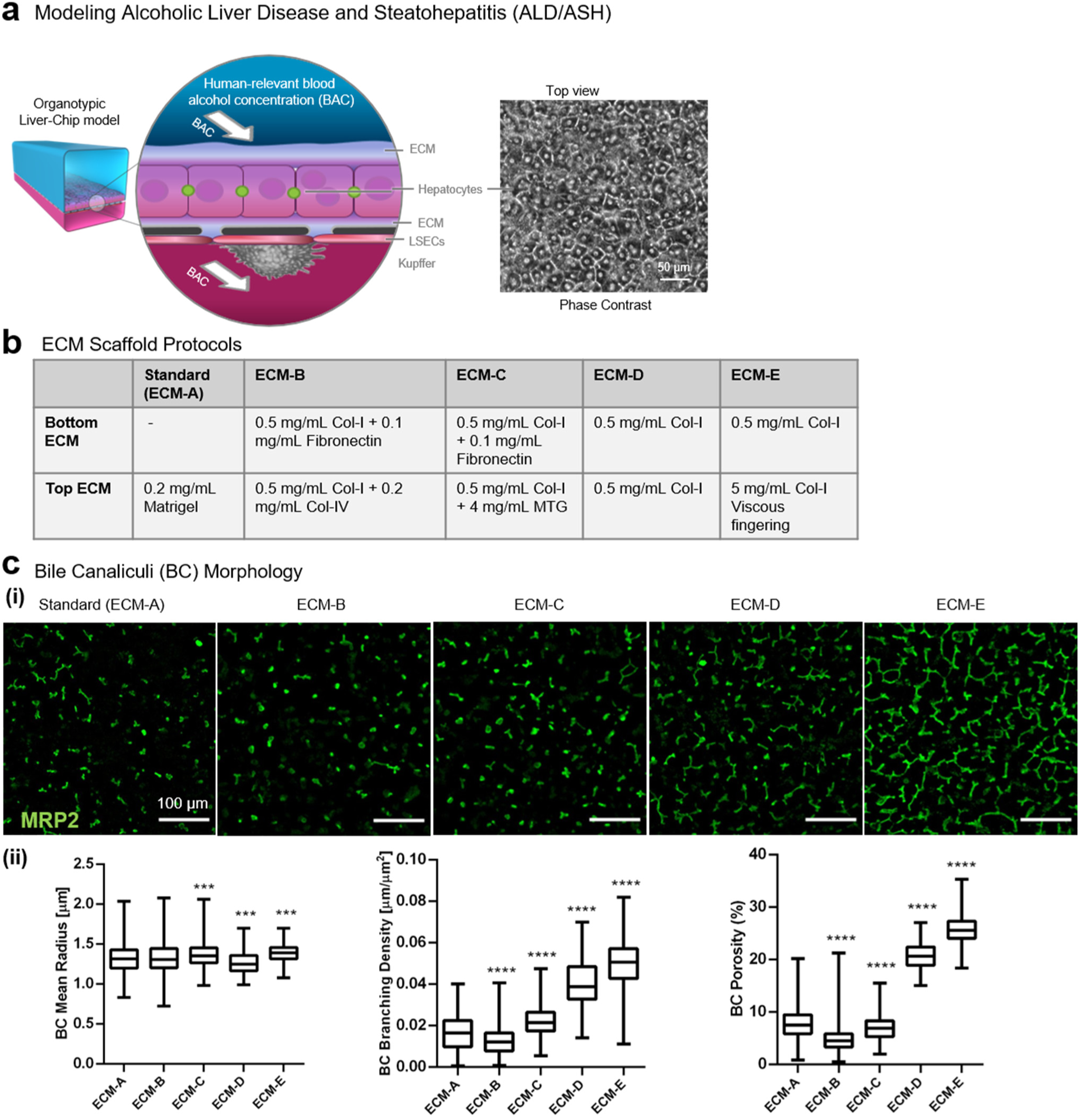
Development of the ALD Liver-Chip. **a**. Approach for modeling human ALD/ASH by exposing the organotypic Liver-Chip to human relevant blood-alcohol concentrations (BAC). **b**. Details of experimental ECM scaffolding protocols tested in the Liver-Chip. **c**. Optimization of BC network integrity. **(i)** Representative images showing vastly different BC networks (MRP2, green) of the tri-culture Liver-Chip under different ECM conditions. **(ii)** Effects of different ECM conditions on the radius, branching density, and area fraction (tissue porosity) of BC networks as assessed by digital pathology. Biomimetic hepatobiliary architecture (condition ECM-E) is characterized by higher branching density, higher porosity and more narrowly distributed mean radius compared to control (condition ECM-A). Data are from one experiment with n=3 chips per condition. Data represent median ± (min and max), \**p*< 0.05; \*\**p*< 0.01, \*\*\*\**p*< 0.0001 versus control (Branching density and porosity: Kruskal-Wallis and Dunnett’s multiple comparisons test; Radius: KS Test and Bonferroni correction for multiple comparisons test).

### Ethanol-induced steatosis in the Liver-Chip

Hepatic intracellular lipid accumulation, or steatosis, is the key histological finding of FLD caused either by ethanol abuse or high-fat diet^10,14^. Therefore, to confirm that the Liver-Chip is suitable for modeling ALD and its progression to ASH, we first probed the capability of the Liver Chip to demonstrate lipid accumulation in hepatocytes on chip upon treatment with ethanol. As a positive control for induction of steatosis, we used fatty acids, a reproducible experimental method for induction of a steatotic phenotype^31^. We monitored the hepatic lipid accumulation using digital pathology to automatically segment images of AdipoRed-stained tissues and quantify the lipid droplets frequency and size. Ethanol concentrations ranging from 0.08% to 0.16% (i.e., 80 mg/ml to 160 mg/ml, or 17.4mM to 34mM) were chosen to mimic the alcohol concentrations found in the blood (BAC) of human patients after alcohol consumption^32^. For comparison, a BAC of 0.08% is the upper limit to for legal driving in the United States and the UK. In order to demonstrate the sensitivity of the Liver Chip to ethanol in a reproducible manner, we had to adjust the medium the hepatocytes were exposed during the treatment, for glucose, insulin and cortisol concentration, as described in Methods. Steatosis was induced following treatment with either ethanol, at BACs ranging from 0.08% to 0.16%, or with oleic acid (1 µg/ml) for 48hrs (Fig 2a,i). While triglyceride (TG) storage did not significantly change over this period of time (Data not shown), likely due to the short treatment window, The average lipid droplet size visualized using AdipoRed staining increased with rising ethanol concentrations ^33^,^,34^ (Fig. 2a,ii), indicating the sensitivity of the Liver Chip to operate within a range of clinically relevant BACs, as required for disease modeling.

**Figure 2.**
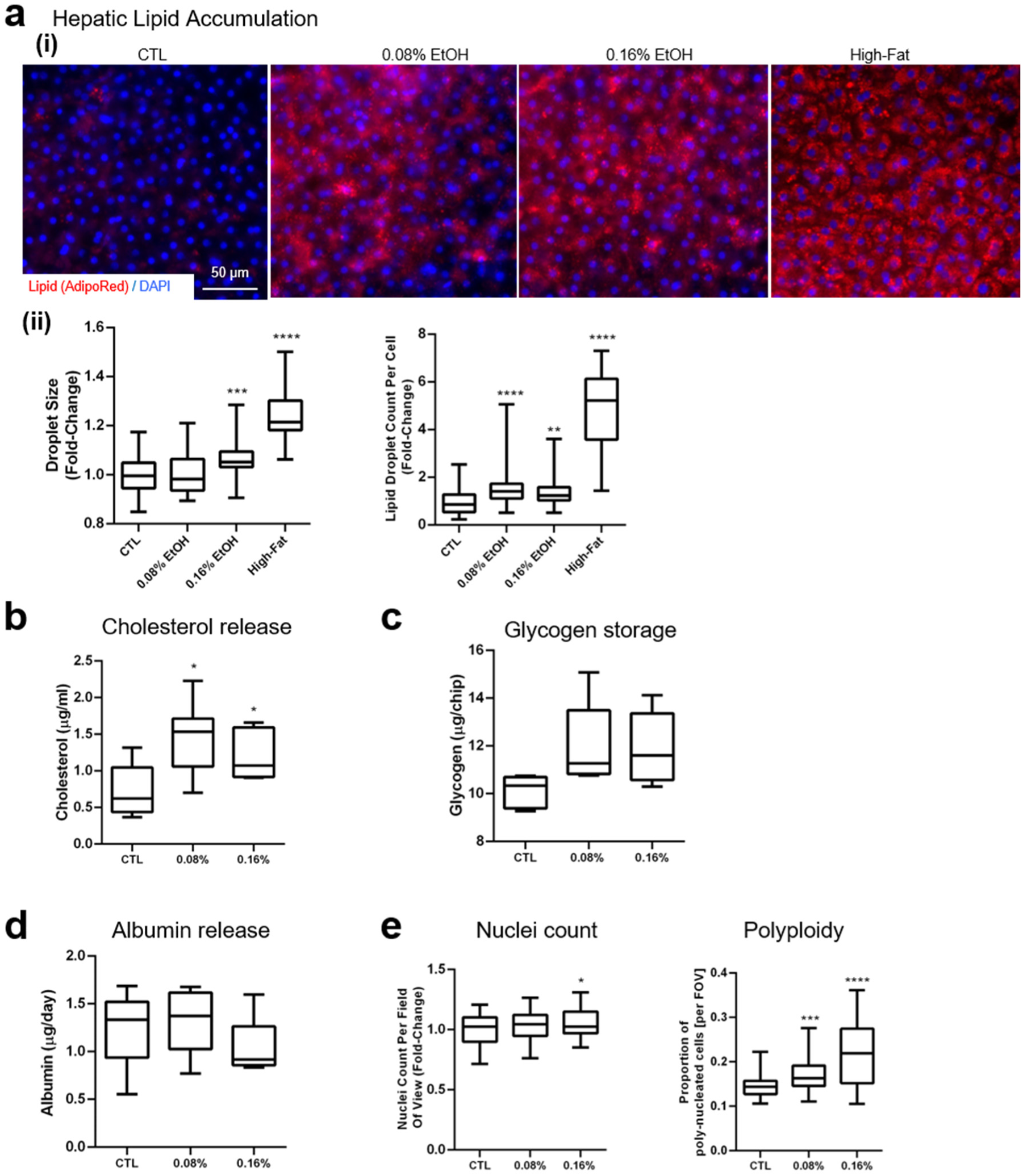
Assessment of liver toxicity, metabolic changes, and polyploidy in the ALD/ASH Liver-Chip. **a**. Hepatic lipid accumulation. **(i)** Lipid droplet accumulation in hepatocytes visualized using AdipoRed staining after administration of fat (oleic acid 1 µg/ml; positive control) or ethanol (0.08% and 0.16%) for 48 hours. **(ii)** Digital pathology was used to quantify the number of lipid droplets per cell, and lipid droplet size (projected area). Data represent median ± (min and max). \**p* < 0.05; \*\**p*< 0.01 versus control (Kruskal-Wallis and Dunnett’s multiple comparisons test). **b-d**. Quantitative analysis of hepatic functional markers in the Liver-Chip after 48h of exposure to physiologically relevant BAC. Fluorometric assessment using ELISA of (b) cholesterol levels in the effluent, (c) glycogen storage in cell lysate, and (d) albumin release. **e**. Nuclei count per field of view and proportion of hepatocytes with multiple nuclei per cell (polyploidy).

Next, we assessed whether the lipid accumulation observed in the Liver-Chip in response to ethanol treatment, was associated with metabolic dysregulation, although this usually requires more prolonged exposure to the challenge^35^. To this purpose we used fluorometric assays for biochemical markers of liver function, including albumin, cholesterol, glucose, and glycogen (Fig. 2b-e). We found that the 48hrs exposure to ethanol at human-relevant BACs resulted in a significant increase in cholesterol levels in the effluent (Fig. 2b), and to a lesser degree, in the cell lysates (Supplementary Fig 2A). We confirmed the reproducibility of this finding in Liver-Chips populated with hepatocytes from 2 different donors, that showed that despite the donor-to-donor variability in the baseline levels (Supplementary Fig. 3A), there was no difference in the cholesterol release in response to ethanol between the two donors (Supplementary Fig. 3C). Glycogen storage, as determined in the cell lysates, showed a trend for increase upon exposure to 0.08% ethanol (Fig. 2c), while no changes were detected in glucose release (Supplementary Fig. 2B), except when we used toxic ethanol concentrations above 0.32% (data not shown). Similarly, albumin release was not affected by this, of limited length, exposure to ethanol (Fig. 2d), in line with clinical data in patients where such changes develop over time, and characterize a moderate to more severe deterioration of liver function^36,37^. Interestingly, while exposure to ethanol had a minimal effect on the number of hepatocytes nuclei counted per field of view, use of image processing to automatically segment the cellular boundaries and determine the number of nuclei per cell, revealed ethanol-induced increases in polyploidy, thought as a marker of tissue stress ^38^ (Fig. 2e).

**Figure 3.**
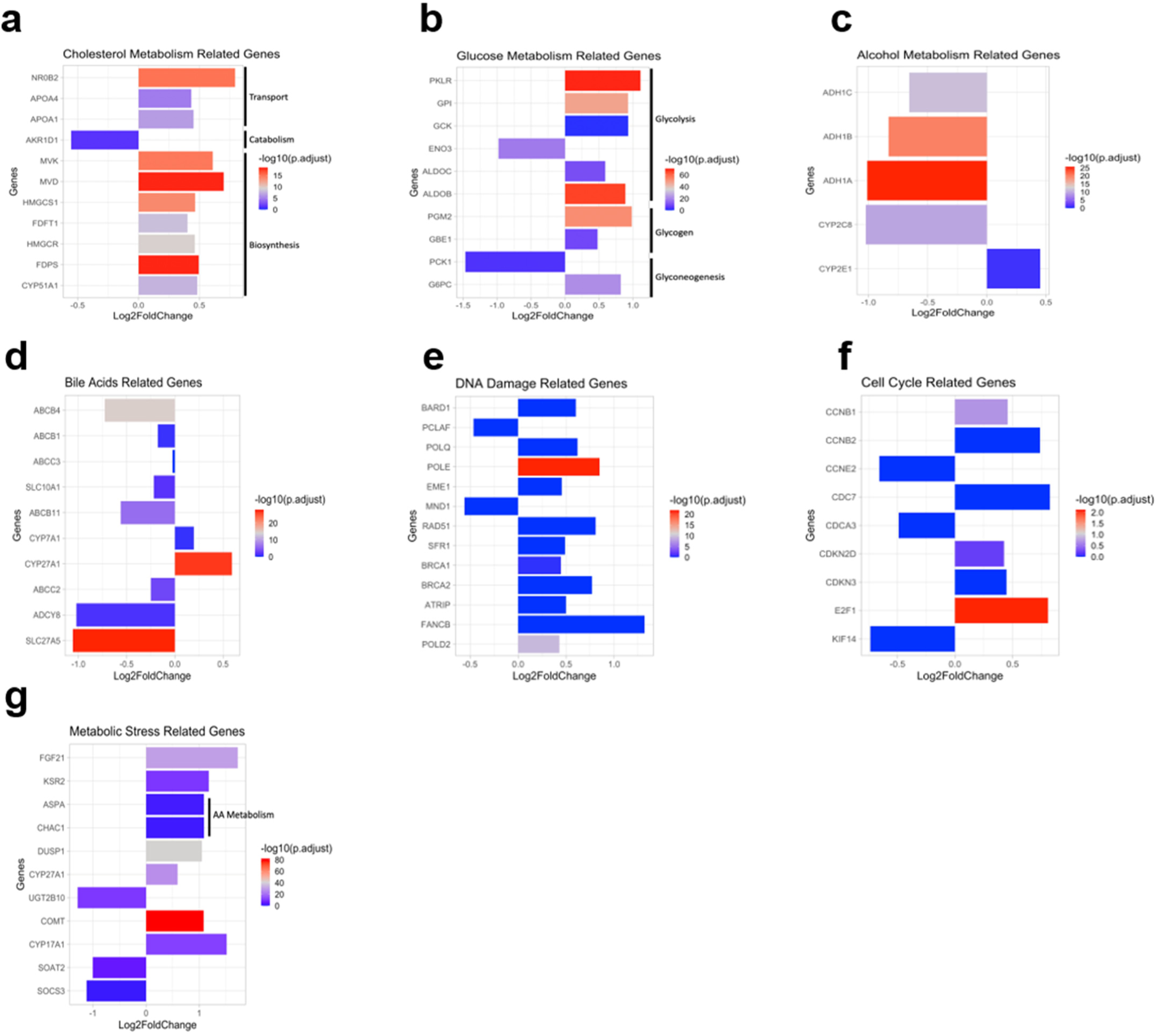
Gene expression profiling of the Liver-Chip induced by physiologically - relevant ethanol concentrations. Differential gene expression analysis in hepatocytes from ethanol-treated (exposed for 48h at ethanol concentrations of either 0.08% or 0.16%, see Methods) and control Liver-Chips revealed significant differences in the expression of genes related to **a**. alcohol metabolism, **b**. cholesterol metabolism, **c**. glucose metabolism, **d**. bile acid production and maintenance (i.e., cholestasis), **e**. DNA damage, **f**. cell cycle regulation, and **g**. oxidative and metabolic stress. Adjusted p-values are demonstrated by bar color. Data from one experiment with 2-5 chips per condition.

To assess for changes in gene expression underpinning the above phenotypic responses to ethanol exposure, we performed RNA-seq analysis. The differentially expressed (DE) genes and the associated stratification, based on the magnitude of differences between ethanol-exposed and control chips, were plotted as shown in Supplementary Fig. 4A. In the group of the ethanol-exposed chips (at concentrations of either 0.08% or 0.16%), we identified 123 DE genes (red dots, adj. p-value<0.05 and |log_2_FoldChange|>1), of which 87 were upregulated (depicted in red in the heatmap) and 36 were downregulated (depicted in blue in the heatmap) (Supplementary Fig. 4B). Pathway analysis of the DE genes using the KEGG bioinformatics database revealed changes in the expression of genes implicated in functions consistent with the phenotypic and other readouts discussed above (Supplementary Fig. 4C). Relevant to lipid metabolism, we found upregulation of three genes related to cholesterol transport (*NR0B2*^39^, *APOA4*^40^, and *APOA1*^41^) and another seven genes related to cholesterol biosynthesis (*MVK, MVD, HMGCS1*^42^, *FDFT1*^43^, *HMGCR*^44^, *FDPS*^45^, *CYP51A1*^46^), along with downregulation of *AKR1D1*^47^, involved in cholesterol catabolism (Fig. 3a). Specifically, glycolysis/gluconeogenesis, steroid hormone biosynthesis, and insulin resistance/type II diabetes were among the key pathways affected by exposure of the Liver-Chip to ethanol. Further, the ALD Liver-Chip, as depicted in the altered expression of genes involved in glycolysis, gluconeogenesis, and glycogen metabolism (Fig. 3b). Ethanol-treated Liver-Chips also exhibited significant changes in the expression of members of the alcohol dehydrogenase (ADH) gene family, including *ADH1C*^48^, *ADH1B*^49^ and *ADH1A*, all involved in alcohol metabolism ^49^.Most importantly,, we found that alcohol induced the expression of CYP2E1, critical for ethanol oxidation and the associated induction in oxidative stress (Fig. 3c). We detected similar effects in genes associated with induction of oxidative stress, a major pathogenic mechanism of ALD^50^. Further, exposure to ethanol affected the expression of clinically-relevant genes involved in bile acid production and processing, hallmarks of cholestasis. Here, we identified changes in the expression of a number of specific genes such as the major BC transporters *ABCB4 (MDR3)* ^51^, *ABCB1 (MDR1)*^52^, *ABCC3 (MRP-3)*^53^, *SLC10A1 (NTCP)*^54^, *ABCB11 (BSEP)*^55^, *ABCC2 (MRP-2)*^55^, *and ADCY8*^56^ or genes associated with bile acid processing such as *SLC27A5*, involved in fatty acid elongation^57^, *CYP7A1*^58,59^ which regulates the overall bile acid production rate and *CYP27A1*^60^, associated with the bile acids synthesis alternative pathway (Fig. 3d). Our data also show that the ALD Liver-Chip picked up the impact of ethanol treatment on genes involved in the metabolism of alanine, aspartate, and glutamate. In line with previous data shown the link between FLD and DNA damage in hepatocytes^18^, our analysis revealed upregulation of *POLE* and *POLD2*^61^, both involved in DNA replication and repair, as well as of *RAD51* and *FANCB*, which are expressed at DNA damage sites and implicated in homologous recombination^62,63^ (Fig. 3e). Further, we saw upregulation of *E2F1*^64^ and *CCNB1*^65^ and downregulation of *KIF14*^66^ and *CCNE2*^67^, all participating in cell cycle regulation (Fig. 3f). Notably, several of the DNA damage-associated genes identified in our data have not yet been implicated in the pathogenesis of FLD pathogenesis, suggestive of the potential of the ALD Liver-Chip platform to be applied for detailed characterization of the alcohol-induced DNA damage. Lastly, ethanol exposure led to the dysregulation of several markers of oxidative stress in the Liver-Chip, such as genes of the metallothionein family (*MT1)*^68–70^ (Supplementary Fig. 4C) (Fig. 3g) andaltered the expression of *DUSP1*^71^ (Fig. 3g), in further support of the stress response induced by ethanol in the liver chip.

**Figure 4:**
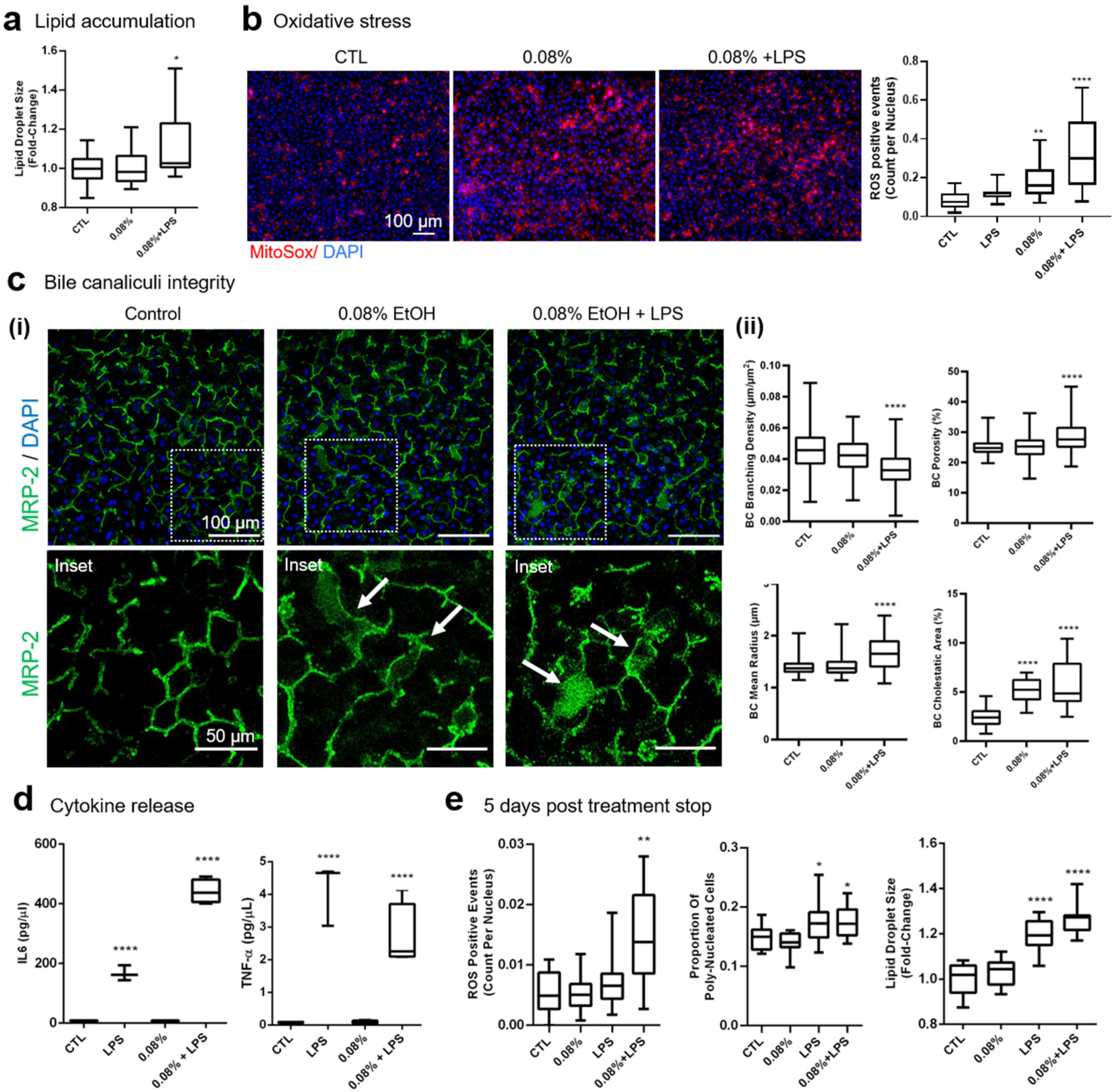
Modeling two-hit hypothesis for ALD/ASH and alcohol abstinence in the ALD Liver-chip. Data was collected following 48hrs of exposure to relevant BAC 0.08%ethanol) or ethanol+LPS. **a**. Quantification of mean lipid droplet size in the Liver-Chip hepatocytes. **b**. Representative images of MitoSox staining in the Liver-Chip hepatocytes (left) and quantification of ROS events (right). **c**. Cholestatic responses to treatment. **(i)** Representative images showing changes in MRP-2 bile canaliculi staining (green) in the Liver-Chip (condition ECM-D). **(ii)** Quantification of the changes in bile canaliculi radius, branching density, and porosity, as well as cholestatic area in the Liver-Chip. **d**. Release of IL-6, and TNF-alpha as measured by multiplexed immunoassays. Data from 2 (48h ethanol) or 1 (48h high fat diet; 5 days recovery) independent experiments, minimally n=2 chips per condition, 5-8 images per chip where applicable. Data represent median ± (min and max). \**p*< 0.05; \*\**p*< 0.01, \*\*\*\**p*< 0.0001 versus control (Kruskal-Wallis and Dunnett’s multiple comparisons test). **e**. Quantification of mean lipid droplet size, frequency of oxidative stress events and polyploidy in hepatocytes after 48h of exposure of the Liver-Chip to either ethanol or ethanol + LPS followed by 5 days of recovery following cessation of exposure to ethanol.

### Modeling two-hit hypothesis for ALD/ASH development

Having established that the Liver-Chip responds to ethanol, we next explored the possibility of modeling the progression of ALD. According to the two-or second-hit hypothesis, simple steatosis induced by alcohol requires a second insult for progression to ASH^72^. Ethanol consumption compromises the intestinal barrier function resulting in increased permeability to intestinal endotoxins, like bacterial lipopolysaccharide (LPS), from the gut to the liver via the portal vein^20^. The leaky barrier is thought to act as the critical second hit in the progression of ALD to inflammation (ASH)^73^. Therefore, we tested whether co-exposure to ethanol and LPS would worsen steatosis and the oxidative stress in the ALD Liver-Chip. Using AdipoRed staining, we found that the size of hepatic lipid droplets was increased in Liver-Chips exposed for 48hrs to 0.08% ethanol+LPS, whereas treatment with ethanol alone at this concentration did not have an effect (Fig. 4a). To asses for induction of oxidative stress, we used MitoSox staining, a fluorescent marker of mitochondrial reactive oxygen species (ROS) and quantified the number of ROS events per nuclei. Our findings revealed a dose-dependent increase in ROS events in response to ethanol, which, indeed, was further exacerbated upon co-exposure to ethanol and LPS. The specificity of the response is highlighted by lack of any effect upon treatment with LPS alone (Fig. 4b). These results are in line with our RNAseq data showing the effects of alcohol on genes related to DNA damage and cell cycle and are consistent with the reported disease mechanisms. Induction of CYP450 by ethanol and free fatty acids and the consequential oxidative stress rampage have previously been implicated as critical mechanisms for the increased DNA damage in hepatocytes and associated progress to hepatocellular carcinoma (HCC)^18^. However, this is the first, human relevant, *in vitro* model that can pick up oxidative stress and DNA damage induction by ethanol or LPS and ethanol together, possibly due to the sustained expression of P450, such as the ALD-relevant CYP2E1.

Inflammatory cytokines are significantly elevated in alcoholic patients with advanced liver disease^74^ and have been proposed as therapeutic targets, as inhibition of TNF-α action was protective against alcohol-induced liver injury in mouse models^75^. Therefore we assessed the secretion of cytokines in response to ethanol and LPS, as our Liver-Chip model contains Kupffer cells which are a major source of proinflammatory cytokines^76^. Whereas as anticipated treatment of the Liver-Chip with LPS robustly increased the release of the proinflammatory cytokines IL-6 and TNF-α co-treatment with 0.08% ethanol + LPS resulted in further increase in the production of IL-6 (Fig. 4d), similar to findings from clinical and experimental studies demonstrating the role of IL-6 in the progress of the disease ^74,77,78^. The cytokines rise is most likely linked to the increase in oxidative stress (Fig. 4b) and the associated cell injury or other mechanisms as previously described^79^.

Patients with alcoholic liver disease frequently manifest clinical and/or histologic evidence of cholestasis^80^, a term defining as either decrease in bile flow due to impaired secretion by hepatocytes or obstruction of bile flow through the intra-or extrahepatic bile transport network. Cholestasis is diagnosed based on accumulation of the potentially toxic cholephiles in the liver and the systemic circulation^80^. Currently, the mechanism of alcohol-induced cholestasis remains poorly understood. We have recently shown the ability of our human Liver-Chip to develop bile canaliculi and recapitulate in vivo findings on drug-induced destruction of bile canaliculi transporters^19^. In the current study, the Liver-Chip was optimized to develop more extensive, biomimetic BC networks. Given that RNA seq analysis of hepatocytes in the alcohol-treated Liver-Chip depicted altered expression of bile canaliculi-related genes, we used quantitative analysis of the MRP-2 stained BC network to assess the sensitivity of the Liver-Chip and demonstrated alcohol-induced cholestatic changes at the level of the canaliculi (Fig. 4c,i). In the Liver-Chips treated with 0.08% ethanol, we identified areas of markedly dilated BCs, a marker of cholestasis, that were significantly expanded in those with 0.08% ethanol + LPS together, as anticipated by the more severe hepatic injury (Fig. 4c,ii). Further, in the latter the average BC radius increased throughout the BC network, whereas the branching density was decreased, both signs of the worsening of hepatic damage, in line with the two-hit hypothesis for the pathogenesis of ASH/NASH^22^.

We would like to also mention here our preliminary study on leveraging the optimized ECM scaffold (Fig. 1b and Supplementary Fig. 1C) for embedding of hepatic stellate cells (HSCs). This approach enabled the development of a quad-culture Liver-Chip (Supplementary Fig. 5A) used for a proof-of-concept experiment to evaluate the contribution of HSCs to the onset and progression of ALD in the Liver-Chip. Our preliminary results suggest that HSCs added to the Liver-Chip robustly embedded in the 3D matrix (Supplementary Fig. 5B and C) and supported a quad-culture Liver-Chip, which showed more pronounced ethanol concentration-dependent steatosis (Supplementary Fig. 5D) than the tri-culture Liver-Chip (no HSCs) (Supplementary Fig. 5E).

### Modeling of the hepatocyte recovery following abstinence from alcohol with the Liver-Chip

Clinical data have shown the potential of the steatotic liver to repair the associated hepatocyte injury following timely abstinence from alcohol or with diet modifications^21^. Thus, we assessed the responses of the “alcoholic” Liver-Chip to withdrawal from ethanol or ethanol + LPS. Treatment of the Liver-Chip for 48h with ethanol or ethanol + LPS as above was followed by a 5 day-long treatment-free “recovery” period. By the end of the recovery period, ethanol withdrawal resulted in normalization of oxidative stress whereas this was not the case for the Liver-Chips exposed to ethanol+LPS together. Polyploidy and lipid droplet size also normalized in all treatments that did not involve LPS (Fig. 4e). Although it is possible that a longer recovery period might lead to recovery of all treatment groups, these data are promising as they suggest that the ALD Liver-Chip shows an insult-dependent recovery, which parallels the lack of repair of hepatic injury in patients with more advanced alcoholic disease. Further, they underscore the potential of the Liver-Chip to model recovery from ethanol-induced damage in a clinically relevant way, and to uncover potential new targets to leverage regenerative mechanisms in the human liver for therapeutic or prognostic purposes.

## Discussion

Considering the global rise in alcohol-induced liver disease and associated co-morbidities, there is a great need to advance human in vitro models to understand disease mechanisms and support the development of safe and efficacious diagnostic and therapeutic approaches for FLD. Here we describe how we leveraged and further optimized our recently developed human Liver-Chip platform^76^ to address an unmet medical need, the preclinical modeling of progressive alcohol-induced liver disease (ALD). The human Liver-Chip supports a complex co-culture system containing up to four of the main cell types found in the liver and recapitulates relevant cell-cell interactions, improves nutrients availability in the cells via continuous perfusion and maintains long-term cell viability. The unique properties of this design allow for continuous exposure to ethanol followed by withdrawal in the same sample for modeling recovery. It also enables the study of several ALD endpoints and multi-modal assessment of the injury in the same chip. We have recently shown the use of our human Liver-Chip platform for creating safety profiles and studying toxicity for drugs in development and confirm clinically relevant mechanisms of action^19^. In this study we describe how we augmented the human Liver-Chip system with new ECM designs and quantitative readouts to model aspects of alcohol-induced liver disease and assess them with clinically relevant endpoints.

We demonstrate how perfusion of the Liver-Chip with relevant blood alcohol concentrations (BAC) to model heavy drinking resulted in pathological changes associated with ALD in a sensitive and reproducible manner. As our primary goal was to leverage the Liver-Chip that was previously optimized for safety and toxicity studies^19^ and, by adding new capabilities, advance it as a platform for modeling human liver diseases, we applied a short-term experimental design to conduct proof-of-concept studies. We show that the ALD Liver-Chip model recapitulated disease endpoints such as intracellular accumulation of lipids, development of oxidative stress, and cholesterol synthesis dysregulation upon exposure to alcohol for 48hrs^21,81^. RNA seq for gene expression profiling supported all the experimental findings and hinted at additional disease-relevant pathways. Importantly, key alcohol-induced changes were reversed by withdrawal of ethanol for several days, mirroring the effects of alcohol abstinence in human patients. Furthermore, we introduced innovations to the Liver-Chip model that greatly increases its range of applications for translational studies. By optimizing the chemical and mechanical properties of the ECM patterning for the Liver-Chip we induced a stable, biomimetic bile canaliculi network in the hepatic tissue, as assessed by quantitative imaging metrics. Of note, *in vitro* human hepatocyte cultures have thus far failed to demonstrate sustained formation of BC networks. Our advances hence augmented the human Liver-Chip with new capabilities by enabling visualization and quantification of changes to the biomimetic BC network, a sensitive maker of liver injury^22,23^, which was also confirmed in our study. We also applied the newly optimized ECM protocol in a proof-of concept experiment to include human hepatic stellate cells in the Liver-Chip. We report here our promising early findings on the potential impact of HSCs on promoting ethanol-induced steatosis, as modeled in the ALD Liver-Chip.

We have previously shown that the expression and functionality of key enzymes of the CYP450 family, such as the ALD-relevant CYP2E1A, are maintained in this platform for prolonged periods, as compared to most of the currently used hepatocyte culture systems^19^. The ALD Liver-Chip exhibits alcohol treatment-induced effects in the expression of genes related to nutrient metabolism and metabolic stress, which is in agreement with the dysregulation of steatogenic enzymes and transcription factors identified by in vivo studies^9^,such as CYP2E1A, HMGCR and SCL27A5^82^. We also identified ethanol-dependent dysregulation of cell cycle- and DNA damage-related genes. Furthermore, the ALD Liver-Chip exhibited altered gene expression of cholesterol metabolism and BC transporters, a finding corroborated by reduced BC network integrity as assessed by quantitative imaging. Thus, our Liver-Chip provides a very promising system to model human ALD and study clinically relevant metabolic events, such as ethanol metabolism, lipogenesis, biliary function, and oxidative stress.

Next, we assessed the capability of the Liver-Chip to simulate progression of ALD, for instance when liver injury is combined with leaky gut. Dysbiosis is a chronic alcohol-driven intestinal complication that leads to increased permeability of the intestinal lining and systemic release of intestinal endotoxins from the gut into the systemic circulation. The circulating endotoxins drive hepatic inflammation and release of proinflammatory factors which further induce tissue damage and deterioration of liver function through activation of Kupffer cells, and Kupffer cell-derived inflammatory cytokines, such as TNF and IL-1^75,83^. Here, we modeled this scenario by perfusing the Liver-Chip with LPS and ethanol combined. We show that this treatment increased oxidative stress (as per ROS quantification) and hepatocyte polyploidy, a suggested liver response to stress that is protective against hepatic carcinoma^38^, and it compromised the integrity of the BC network. Moreover, withdrawal from LPS and ethanol for five days did not diminish the oxidative stress or polyploidy, in contrast to the rescuing of the pathology induced by ethanol alone. This suggests that chronic alcohol intake coupled with systemic inflammation worsens the liver damage and may significantly compromise recovery in ALD/ASH, similar to the responses of patients with more advanced disease^84^. In animal experiments administration of antibiotics to reduce endotoxemia or inactivation of Kupffer cells with gadolinium chloride can both prevent liver injury^85^, suggesting that the ALD Liver-Chip model could be useful as a platform to determine human relevancy of proposed mechanisms for new therapeutic approaches.

The ALD Liver-Chip presents multiple advantages over other *in vivo* and i*n vitro* FLD models. Rodent models of NAFLD/NASH, although they typically develop the disease histopathology and have been instrumental in the elucidation of main pathogenetic mechanisms such as insulin resistance, they were not successful in recapitulating the variability in the patients responses^86–88^. With respect to rodent models of ALD/ASH, unfortunately they have been notoriously resistant to the hepatotoxic effects of alcohol alone, and they only develop significant chronic liver injury when exposed to combinations of alcohol either with a toxin or major dietary modifications^89,90^, and no fibrosis development, which is the most important pathological feature in ALD. Other, non-rodent, animal models such as *Caenorhabditis elegans*, opossum, Ossabaw pig, and primates have been recently introduced, but these studies are still at the early validation stages^87^. *In vitro* models used to further elucidate the molecular mechanisms underlying FLD include standard cultures of primary or immortalized, patient-or rodent-derived, hepatic cells, co-culture and 3D cultures^87^. The main caveats with these models include a lack of the dynamic environment provided in the engineered microphysiological systems and of the in vivo-relevant tissue-tissue interfaces and corresponding cytoarchitecture. As a result, no fluid flow or associated mechanical forces can be applied, oxygen and nutrient transport are limited, and metabolites may accumulate to levels well beyond, or below, the physiologically relevant values, impeding the translation of findings for patients care. In addition, the lack of fluid flow compromises human disease modeling such as the ALD/ASH as it does not support real time perfusion with ethanol to mimic circulating blood alcohol concentrations. Lastly, 3D cultures, arguably the more advanced *in vitro* model, do not support yet visualization of the dynamic changes in cell morphology and cell-cell interactions whereas Organ-Chips do^91^.

In summary, we report how we have combined the human Liver-Chip^19^ with new bioengineering approaches and multimodal profiling to develop an innovative platform for modeling progressive liver injury in response to alcohol (ALD). Our findings demonstrate the potential of the ALD Liver-Chip to model comorbidities and improve translation of preclinical data to the clinic, to uncover novel pathogenic and recovery mechanisms, and to identify the windows for successful intervention in patient cohorts.

## Methods

### Liver-Chip culture

Liver-Chip culture was established according to our previously published protocol^76^. Briefly, primary human hepatocytes (3.5 million cells/mL) from two healthy donors, Donors G and Q, were cultured on one side of a porous membrane (pore size ∼7 microns) in the Emulate microfluidic chip^92^. Extracellular matrix sandwiching and composition in the Liver-Chip was optimized to support BC integrity (Fig. 1b and c, Supplementary Fig. 1 and method sections below). On the other side of the membrane (the lower chamber of the chip) human primary LSECs (3 million cells/mL) and KCs (0.5 million cells/mL) were seeded to mimic the hepatic sinusoid architecture (Fig. 1a). The two cell compartments were perfused independently, and flow rates were optimized to enable optimal survival and maturation of the different cell types^76^. In the Quad Liver-Chip experiments, primary human hepatic stellate cells (50,000 cells/mL) were mixed within the collagen coating (FibriCol at 0.5 mg/ml concentration) and add to activated chips and let incubate overnight before hepatocytes seeding on the top of the collagen and HSC mix. The seeded hepatocytes were overlaid with ECM as described in Fig. 1b and below and maintained in William’s E Medium (WEM) containing Glutamax (Gibco), ITS+ (Corning), dexamethasone (Sigma-Aldrich), ascorbic acid (Sigma-Aldrich), fetal bovine serum (Sigma-Aldrich), and penicillin/streptomycin (Sigma-Aldrich), and incubated at 37°C, 5% CO2. The vascular channel of the Liver-Chip was maintained with human endothelial media (Emulate, Inc.). Two days after seeding, the Liver-Chips were connected to the Human Emulation System(tm) (Emulate, Inc.) and both chip channels were perfused at 30 µL/h to provide a continuous supply of fresh medium for the duration of the experiments. At day 5 in culture, the basal medium was modified to DMEM low glucose (1g/L) supplemented with non-essential amino acids solution (NEAA, 1:200 dilution), glutaMAX (Thermo Fisher) ascorbic acid (Sigma-Aldrich) and penicillin/streptomycin (Sigma-Aldrich). Moreover, ITS+ premix (Corning) was replaced by ITS (GIBCO, 1:500 dilution) in order to achieve a more physiological relevant concentration of insulin, while the FBS supplement was removed.

### Treatment

To model FLD, cell culture medium was supplemented at day 7 in culture either with ethanol (0.04%, 0.08% or 0.16%), with or without lipopolysaccharides (LPS, 1µg/ml), or fatty acid (oleic acid, 1µg/ml). The Liver-Chip was maintained for 48h in treatment medium and either assayed or allowed to recover for 5 days in basal medium.

### Optimization of Liver-Chip ECM for BC integrity

We iteratively modified the original ECM coating protocol^76^ in order to improve BC formation. Five different combinations of ECM composition and deposition strategy were tested (Fig. 1b), where ECM-A denoted the original protocol. Briefly, for ECM-A, top and bottom channels were coated by incubating with 100 µg/mL rat tail collagen-I (Corning) and 25 µ/mL fibronectin (Gibco) overnight at 37°C. Hepatocytes were seeded as described above. After seeding, Matrigel^®^ prepolymer solution was prepared on ice and injected into the top channel, which was then incubated at 37°C overnight. The following day, the top channel was gently flushed with warm medium. In conditions ECM-B, -C, -D, and -E, the hepatocytes are sandwiched between two 3D-ECM gel of different compositions, including Collagen-I (FibriCol^®^), Fibronectin (Gibco), Collagen-IV (Sigma), and the collagen cross-linking agent, microbial transglutaminase (MTG) (Modernist Pantry LLC). The underlying gel was prepared by injecting the gel prepolymer solution after surface activation and incubated overnight at 37°C. The next day, the gel was then flushed twice with 100 µL of warm medium at 187.5 µL/s flow (Eppendorf Xplorer automatic pipet), generating a 3D-ECM on the membrane with a thickness of 30-50 µm (data not shown). Afterwards, hepatocytes were seeded as described above. One day after the hepatocyte seeding, the same method was used to prepare the overlaying gel in conditions ECM-B, -C, and -D. In condition ECM-E, the overlay was prepared via viscous fingering method^30^. Briefly, 5 mg/mL of bovine collagen-I (FibriCol^®^) was injected into the top channel and a pipette tip filled with 200 µL of warm medium was immediately injected into the channel inlet to apply hydrostatic pressure. This will cause an interface instability between medium and prepolymerized collagen-I, enabling the medium to flow through the middle part of the prepolymerized collagen-I, thereby creating a lumen (Supplementary Figure 1C). The chips were then immediately moved into humidified incubator to promote gelation of the lumen formed thick collagen-I ECM on top of the hepatocyte monolayer. ECM-D was used in the experiments in this study, however, ECM-E, which provided additional stability for multilayer matrix-cell scaffolding, was used for the comparison of tri-to quad-culture Liver-Chip.

To assess and confirm ECM scaffold formation during the optimization procedure, we stained the gels with a fluorescent dye: Briefly, 1 mg/mL of N-hydroxysuccinimide (NHS) ester dye (Atto 488 NHS Ester, Sigma Aldrich) was mixed with 50 mM borate buffer (pH 9) in 1:500 ratio. Directly after ECM formation, the prepared staining solution was injected into the top channel and the chips were incubated for 25 min at room temperature in the dark. The top channel was then rinsed three times with PBS prior to fluorescence imaging.

To assess the effect of each ECM protocol on BC network formation, we quantified the MRP-2 stained BC network topology (see method sections on IF staining and Image analysis for details). Optimal conditions thus identified were ECM-D and ECM, and these protocols were then used for engineering ALD/ASH Liver-Chips.

### HSC 2D and 3D culture on plate

To assess the α-smooth muscle actin (αSMA) expression in 2D versus 3D culturing conditions, HSCs were seeded into wells of a 48-well plate either in a 2D-monolayer or in a 3D ECM scaffold (type ECM-D, see Supplementary Figure 1B) that is also used on the Liver-Chip. Briefly, for the 2D culture, HSC suspension (0.05 million cells/mL) was added directly into the wells. For the 3D culture, the HSCs were first suspended in a 0.5 mg/mL Collagen-I (FibriCol^®^) matrix (0.05 million cells/mL) that was then added into the wells. HSC were cultured for 3 days in either conditions, and medium was refreshed every day. After 3 days in culture, the samples were fixed and immunofluorescence staining was performed as described in the separate method section to assess the αSMA activation.

### Immunofluorescence staining

Cells on the Liver-Chips were washed 3x in 1X PBS then fixed with 4 % paraformaldehyde for 20 minutes at room temperature (RT). Chips were then washed 3x with cold 1X PBS and blocked using 1% BSA in 1X PBS for 30 min to 2h at RT. Cells were permeabilized using a 1X PBS solution containing 1% saponin and 10% serum matching the species of the secondary antibody for 30min at RT. Cells were then washed 3x in 1X PBS and blocked again in a 1X PBS solution containing 1% BSA for 2 h to overnight at 4°C. All incubations with primary antibodies were carried out in this blocking buffer overnight at 4°C. This was followed by a two-hour incubation with secondary antibodies (Cell Signaling, Danvers, MA, USA) in the blocking buffer at RT. Immunostaining was performed with the following specific primary antibodies: anti-MRP2 (1:50, Abcam), anti-Vimentin (1:50, Abcam), and anti-smooth muscle actin (anti-αSMA, 1:100, Thermo Fisher). DAPI was used to identify cell nuclei. Images were acquired with either an Olympus fluorescence microscope (IX83) or a Zeiss confocal microscope (LSM880).

### Live cell staining

Liver-Chips were stained in the upper channel with AdipoRed (1:40 dilution in PBS, Lonza) to visualize lipid droplet accumulation, Tetramethylrhodamine, methyl ester (TMRM) (0.1µM in hepatocyte medium, Thermo Fisher) to visualize active mitochondria, and MitoSox^®^ (5uM in hepatocyte medium, Thermo Fisher) and MitoSOX^®^ (5 µM in hepatocyte medium, Thermo Fisher) to visualize cellular oxidative stress, and cholyl-lysyl-fluorescein (CLF, Corning) to visualize bile canaliculi. Each staining solution was prepared and added to the upper channel, incubated for 15-30 min at 37 °C, and washed three times before imaging. NucBlue (Thermo Fisher) staining was used to identify cell nuclei during live imaging. The stained chips were imaged using either an Olympus fluorescence microscope (IX83) or Zeiss confocal microscope (LSM880) and were de-blurred with Olympus cellSens software.

### Image analysis

Analyses of lipid droplet accumulation, ROS events, and nuclei were conducted using ICY^93^, ImageJ-Fiji^94^, CellProfiler^95^, and Matlab (MATLAB, MathWorks Inc., Natick, MA). For ROS measurements, the histogram of the fluorescent images was adjusted to remove the background signal, followed by quantification of ROS events in the region of interest (ROI) based on minimum and maximum size and fluorescent intensity using the batch processing tool in ICY. ImageJ-Fiji was used to preprocess the AdipoRed images for analysis of lipid droplet accumulation. Here, the AdipoRed channel was median filtered, corrected for illumination, and then filtered with a Laplacian filter to emphasize droplet edges and remove larger background structures. The DAPI channel was filtered with an adaptive contrast enhancement algorithm (CLAHE) and then thresholded, followed by multiple dilation and erosion steps to yield a binary image in which nuclei in close proximity, which likely belong to a single poly-nucleated cells, are merged. These preprocessed images were further processed in CellProfiler where a pipeline first automatically segmented the fields of view into estimated cell boundaries using the nuclei as reference points, followed by thresholding and detection of lipid droplets in the AdipoRed channel within each estimated cell boundary. Using CellProfiler modules as well as Matlab scripts, we then computed mean droplet size (i.e., the projected area of the droplet in µm^2^) and the number of droplets per cell. Values of treatment groups were normalized to the median values of the associated control group in order to express fold-change values and thereby mitigate baseline variability due to donor-to-donor and cell batch variability. Furthermore, we computed the proportion of poly-nucleated cells by binning the detected nuclei according to their perimeter, which revealed two distinct populations, i.e. single, well separated nuclei, indicating mononucleated cells, and closely neighboring nuclei fused during thresholding, indicating poly-nucleated cells.

For measuring bile canaliculi network properties, MRP-2 stained chips were imaged at 40x using a confocal point-scanning microscope (Zeiss LSM880, Airyscan). Fields of views were randomly chosen along the entire length of the channel in order to catch heterogeneity caused by ECM deposition or erosion in flow. In each field of view, a z-stack was recorded and combined using maximal intensity projection in order to fully capture the bile canaliculi network. In subsequent image analysis, each field of view was first segmented into 16 sub-windows, and analyzed for three quantitative metrics: Porosity, branching density, and average radius (Supplementary Figure 1C-E). The area fraction taken up by the BC network, which is a measure of porosity, was determined using an ImageJ-Fiji macro that filters and thresholds the signal, followed by the ImageJ particle analysis which detects the area occupied by BC elements (Area BC). BC porosity was then computed as the ratio of Area BC to the total area of the field of view. Average radius and branching density were determined and applying the ImageJ-Fiji plugin “ridge detection”^96^, to detect and measure the radius and length of all BC segments in each window. Then, we computed the average radius measured in each window as well as the branching density, defined as the summed length of the BC branches in the sub-window divided by the area of the sub-window.

The proportion of activated HSCs was determined by counting the proportion of α-SMA positive cells among the vimentin-positive cells found in two chips. For the control group, we found a total of 1 out of 33 vimentin positive cells that were also positive for α-SMA, and for the 0.08% ethanol + LPS treated group that number was 1 out of 47.

### Biochemical assays

#### Cell Lysing

Cells in the Liver-Chip were lysed according to the Protocol for Emulate Organ-Chips (Cell Lysis for Protein Extraction (EP135 v1.0)). In brief, we used Tris lysis buffer (MSD, #R60TX-3]) to directly lyse the cells while still adhering to the chip, collected the lysate, and performed downstream assays.

#### Albumin

Albumin secretion was quantified in Liver-Chip effluent collected from the top channel using the Human Albumin SimpleStep ELISA^®^ Kit (Abcam, #ab179887) according to the manufacturer’s protocol.

#### Cholesterol assessment

Cholesterol was quantified in Liver-Chip effluent according to manufacturer’s protocol for fluorometric detection (Thermo Fisher). Medium in the top channel was changed to standard hepatocyte medium without FBS prior to the experiment. The following sample quantification was used to determine amount of cholesterol in the hepatocytes channel effluent: Net effluent cholesterol = [cholesterol from effluent] µg/ml MINUS [Cholesterol from dosing medium] µg/ml. The same quantification method was used to determine the cholesterol concentration of the hepatocytes cell lysate as described above.

#### Glycogen quantification assay

Hepatocytes in the top channel of the Liver-Chip were lysed as described above, then diluted at a range of 1:500 to 1:1000. Glycogen levels were determined using a standard assay according to manufacturer’s instructions for fluorometric detection (Abcam, #ab65620). **Glucose quantification assay**. Glucose was quantified in Liver-Chip effluent collected from the top channel. Sample concentration were adjusted be within a 50 mg/dl to 200 mg/dl range. Glucose was quantified using a standard kit according to manufacturer’s instruction (Abcam, # ab65333).

#### Gene expression analysis

The cells from both top and bottom channel were separately lysed from the Liver-Chip according to the Protocol for Emulate Organ-Chips: Cell Lysis for RNA Isolation (EP161 v1.0). In brief, we used PureLink RNA Mini Kit lyse buffer (Thermo #12183018) to directly lyse the cells while still adhering to the chip, collected the lysate, and immediately frozen in dry ice. Gene expression levels were analyzed by RNA sequencing performed by GENEWIZ.

#### RNAseq/Pathway analysis

The RNAseq dataset consisted of 2 vehicle samples, 3 samples treated with 0.08% alcohol, and 2 samples treated with 0.16% alcohol. Since differential gene expression analysis between the 0.08% and 0.16% groups yielded no significant differentially expressed genes, we pooled the samples of the two ethanol treatment groups together and constructed a single larger “ethanol-treated” group (consisting of 5 samples) which we compared to the vehicle group.

To remove poor quality adapter sequences and nucleotides, the sequence reads were trimmed using Trimmomatic v.0.36. Next, using the STAR (Spliced Transcripts Alignment to a Reference) aligner v.2.5.2b, we mapped the trimmed reads to the *Homo sapiens* reference genome GRCh38 (available on ENSEMBL). Using the generated BAM files and the feature Counts from the v.1.5.2 subread package, we calculated the unique gene hit counts. Of note, only unique reads that fell within exon regions were counted. We prepared a strand-specific library; therefore, the reads were strand-specifically counted. Using the gene hit counts table, we filtered out genes with very low expression across the samples. The remainder were used for differential gene expression (DGE) analysis. For the DGE analysis, we used the “DESeq2” R package^97^ (Bioconductor) and in order to select the differentially expressed genes, we applied the following thresholds: adjusted p-value<0.05 and |log2FoldChange|>1. Of the 57,500 genes annotated in the genome, 123 were found to have significant differential expression between the vehicle (n=2) and the ethanol exposed chips (n=5). More specifically, 87 (36) genes were found to be significantly upregulated (downregulated) in the ethanol exposed chips (0.08% and 0.16%). These 123 differentially expressed genes were used for the KEGG pathway analysis.

#### Cytokine analysis

Liver-Chip bottom channel (containing Kupffer cells) effluent cytokine levels were measured using the U-PLEX® Biomarker Group 1 Human Assays (MSD^®^ Cat No. K15067L) according to the manufacturer’s instructions.

#### Statistical analysis

As indicated in the figure legends, one-way ANOVA, Sidak’s and Dunnett’s multiple comparisons tests were used for comparing the mean values of parametric data, and the Mann-Whitney U test or Kruskal-Wallis tests followed by Dunnett’s multiple comparisons test was used for comparing the median value of nonparametric data. As noted in figures, the KS-test was occasionally used to test for differences in distributions. All statistical analyses were performed using Prism v6, 7 or 8 (GraphPad). All data was collected from at least 2 independent experiments with at least 3 chips per condition and imaged at 5-10 fields of view per chip (where applicable), unless stated otherwise in the figure legends.

## Acknowledgements

This study was supported in part by an SBIR grant (1R43AA026473-01) from the National Institute of on Alcohol Abuse and Alcoholism and by Emulate. Research reported in this publication was supported by the National Institute on Alcohol Abuse And Alcoholism of the National Institutes of Health under Award Number R43AA026473. The content is solely the responsibility of the authors and does not necessarily represent the official views of the National Institutes of Health. JetPub Scientific Communications LLC, funded by Emulate, provided medical writing support and editorial assistance to the authors during preparation of this manuscript in accordance with Good Publication Practice (GPP3) guidelines.

## Author contributions

DBP and KK developed the experimental design and the overall strategy of the study. JCN developed the strategy and experimental design of the on-chip bile canaliculi optimization. DBP and JCN supervised the experimental studies. KK reviewed all data, data analyses and findings and discussed with GH, DBP, GB, TIM, KS, AW and JCN performed the experiments and collected and analyzed data. JCN developed the quantitative imaging and analysis methods. AS, JCN and TIM performed image analysis. DM conducted the bioinformatics analyses studies. DBP, JCN and DM wrote the manuscript. KK edited and reviewed the manuscript. SL reviewed the manuscript and provided critical points in the discussion and implications of the findings for the field.

## Additional information

**Competing interests:** JCN, DBP, DVM, KK, AS, GH are current or former employees of Emulate and may hold equity interests in Emulate, Inc.

## Supplementary Figures

**Supplementary Figure 1:**
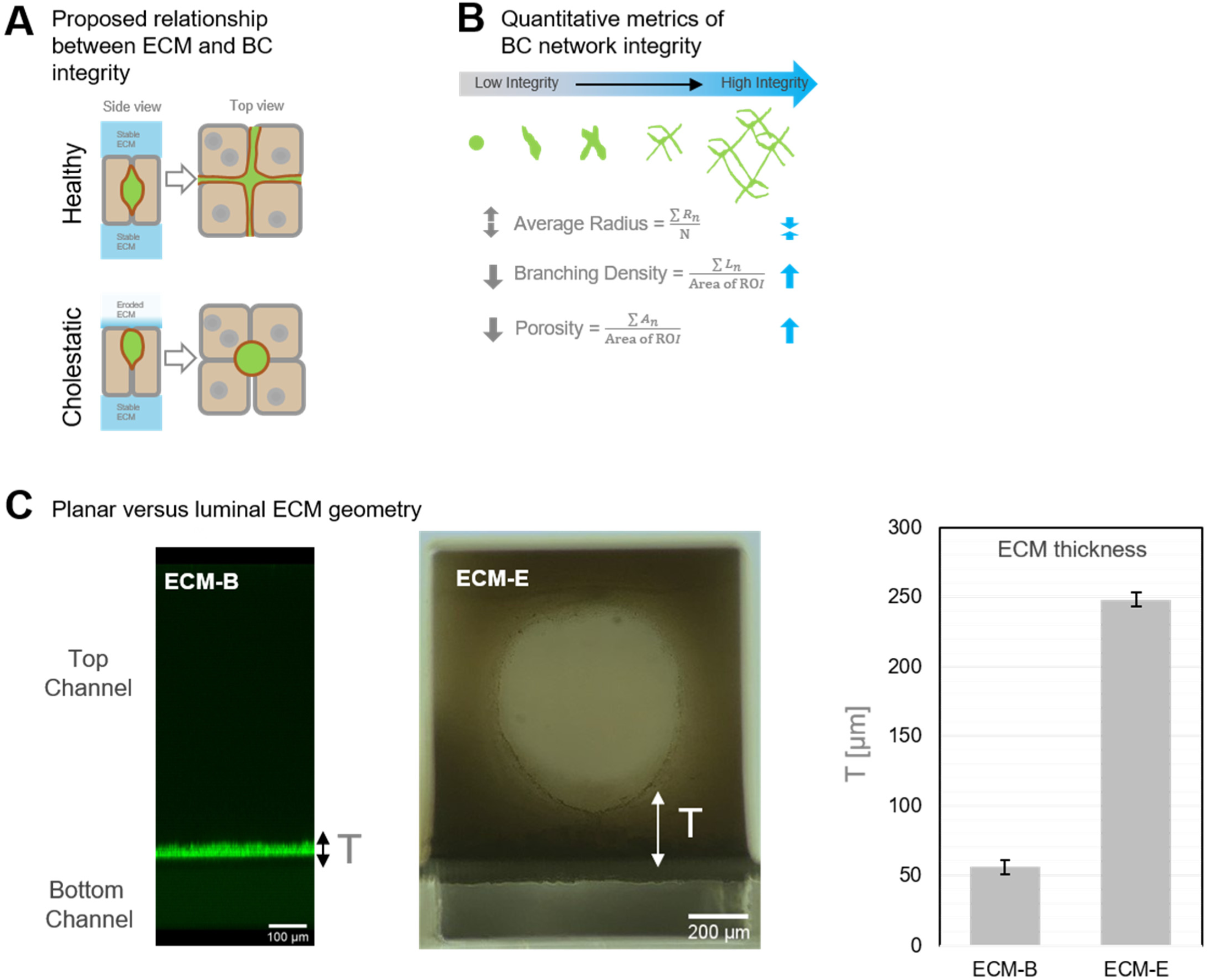
Optimization of bile canaliculi (BC) network integrity. **A**. Schematic of the proposed relationship between the ECM and bile canaliculi integrity based on published work^29^. **B**. Quantitative metrics of bile canaliculi network integrity. BC network integrity is improved when the average radius is narrowly distributed and branching density and porosity increase. **C**. Cross-sectional image and thickness measurements comparing the planar ECM scaffold (here: ECM-B) created by standard membrane coating, and the luminal ECM scaffold (ECM-E) created by viscous fingering. Data are from one experiment with n=3 chips per condition. Data represent mean ± (SEM).

**Supplementary Figure 2:**
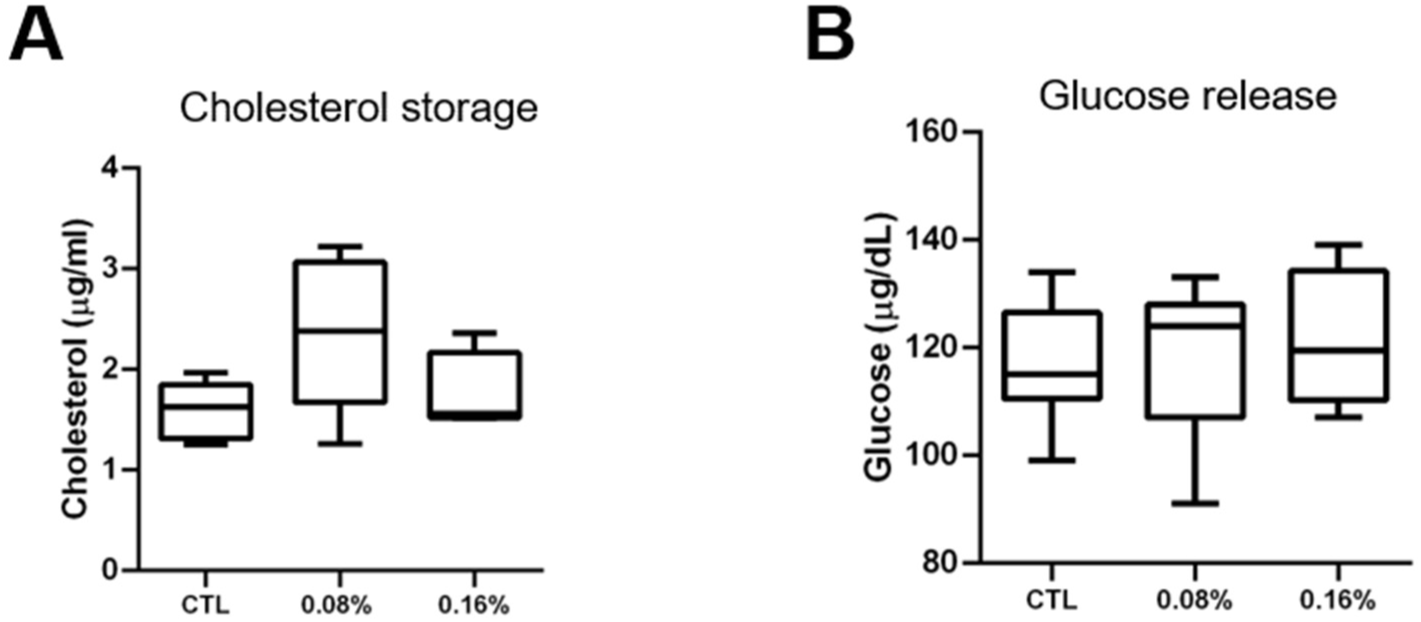
Assessment of metabolic changes in the ALD/ASH Liver-Chip. Data was collected after 48h of exposure to gradually increasing concentrations of 0.08% or 0.16% ethanol. Fluorometric assessment using ELISA of **A** cholesterol levels in cell lysate and **B** glucose release in effluent.

**Supplementary Figure 3:**
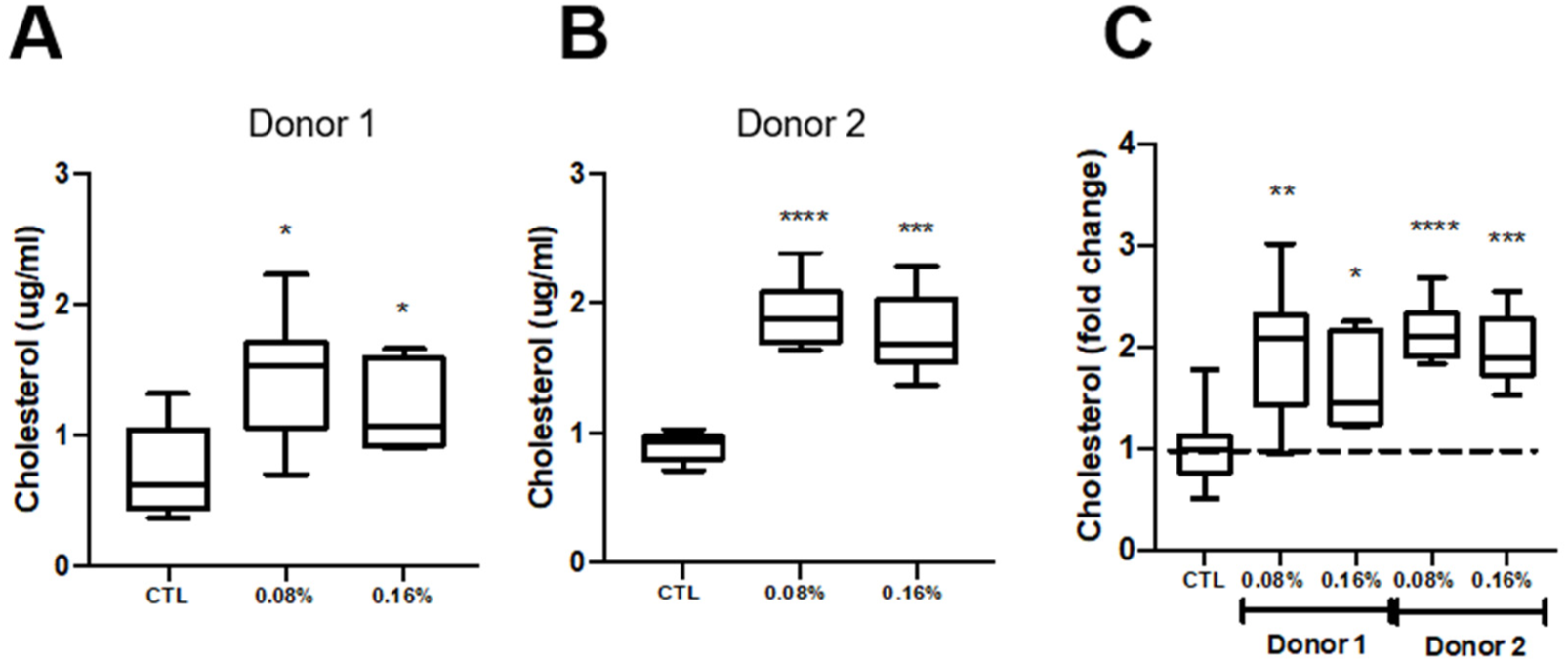
Effect of donor variability on cellular responses to ethanol. **A-B**. Quantification of cholesterol levels in Liver-Chips generated from different donors after 48 h of ethanol exposure; **C**. normalized to show fold-change in cholesterol in response to increasing concentrations of ethanol. Data represent median ± (min and max). \**p*< 0.05; \*\**p*< 0.01; \*\*\**p*< 0.001; \*\*\*\**p*< 0.0001 versus control (Kruskal-Wallis and Dunnett’s multiple comparisons test).

**Supplementary Figure 4.**
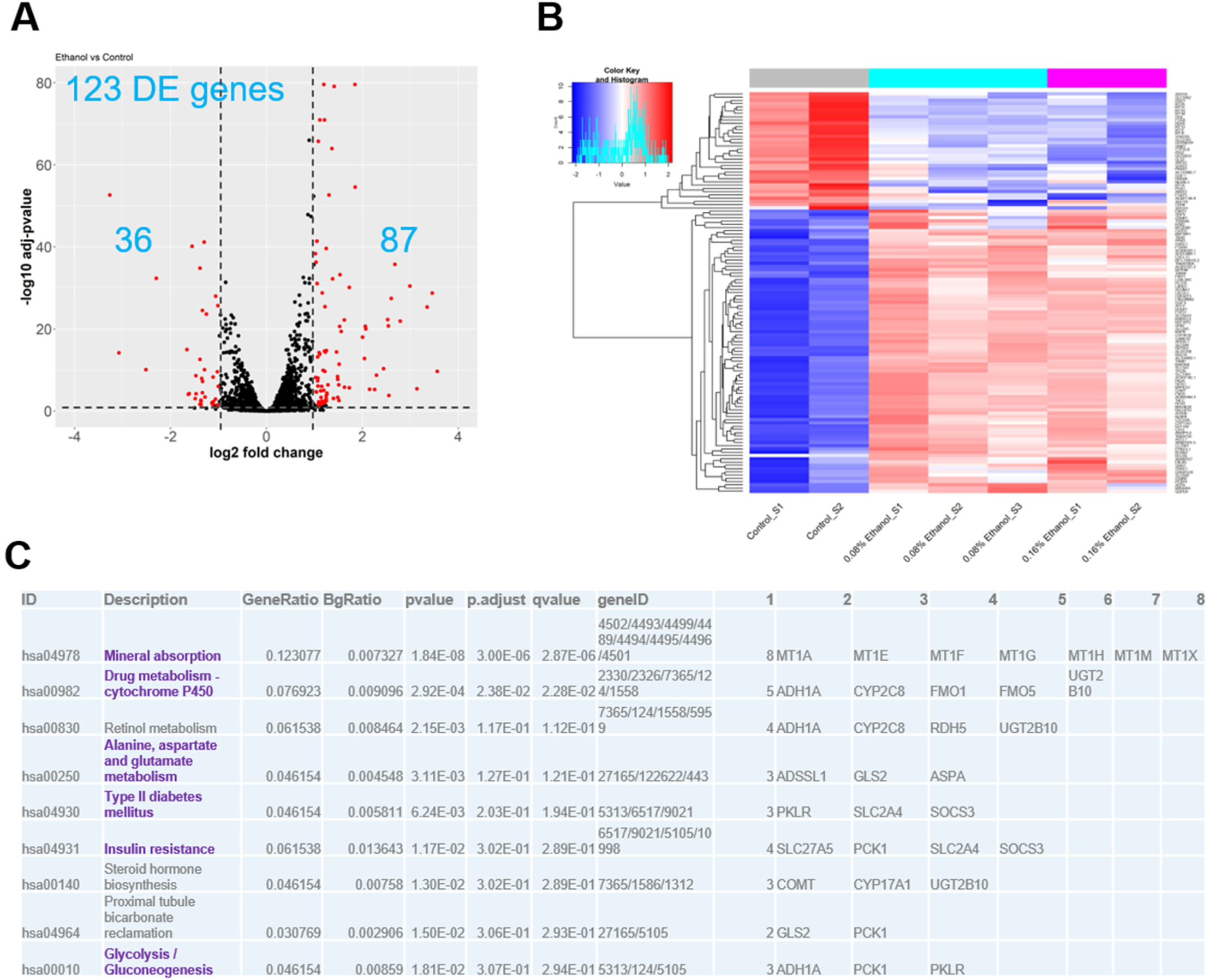
Differential Gene Expression analysis in ethanol-treated vs. control Liver-Chip hepatocytes. Liver-Chips were treated for 48h at ethanol concentrations of either 0.08% or 0.16% (see methods). **A** The volcano plot illustrates the number of differentially expressed (DE) genes and how they stratify based on magnitude of change in ethanol exposed and control Liver-Chips. Red dots: genes that are significantly up-or down-regulated (adj. p-value<0.05 and |log_2_FoldChange|>1); black dots: non-DE genes. **B**. In total, 123 are differentially expressed, 87 are upregulated (red in heatmap) and 36 are downregulated (blue in heatmap) in the ethanol-exposed Liver-chips (0.08% and 0.16%). **C**. Pathway analysis based on the 123 DE genes using the KEGG database.

**Supplementary Figure 5:**
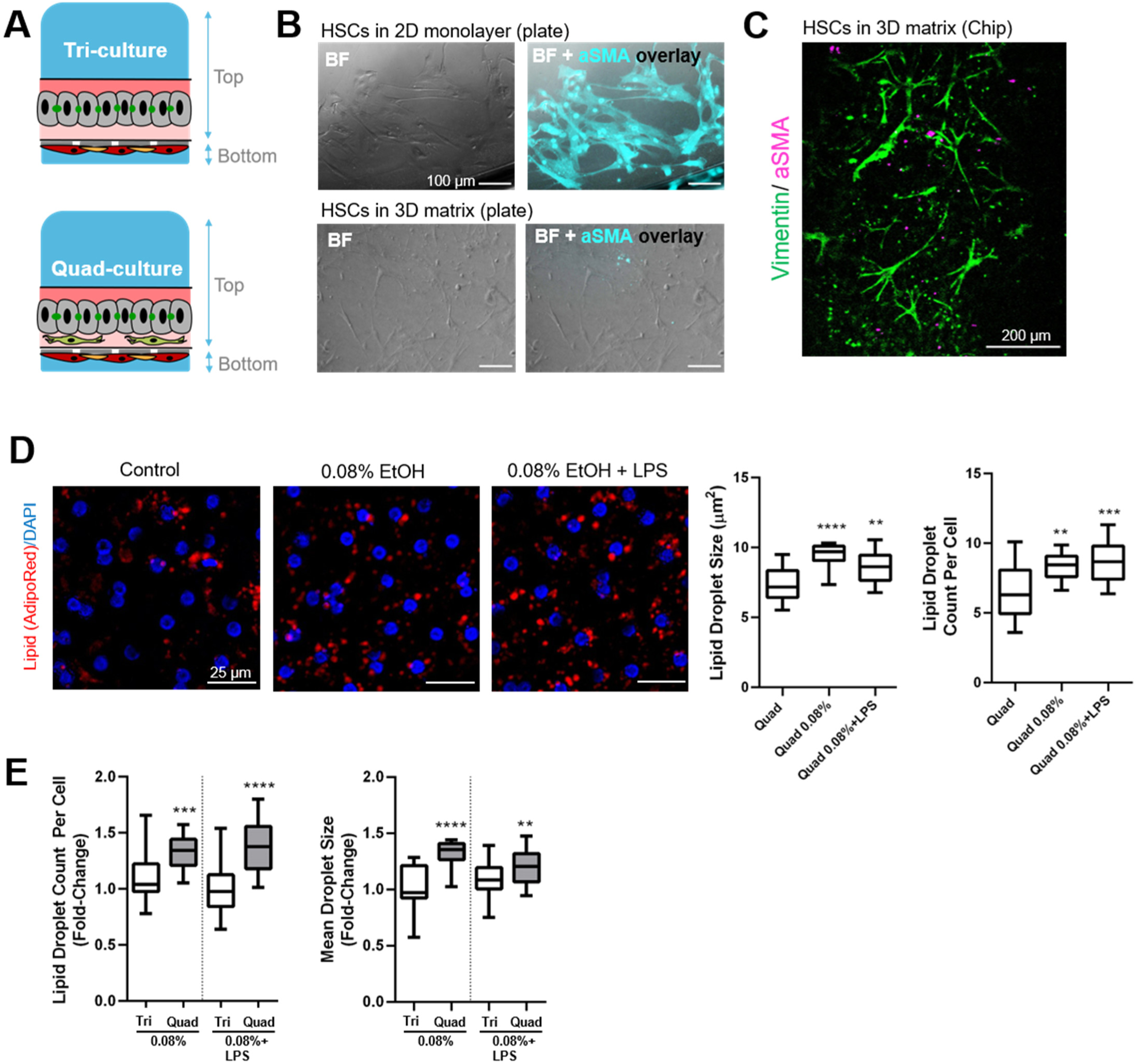
Quad Liver-Chip design and induction of steatosis by ethanol or ethanol + LPS. **A** Schematic illustrating the cellular organization in the quad-culture version of the Liver-Chip containing hepatic stellate cells (HSCs) in the ECM gel bottom layer (condition ECM-E). **B** On-plate test of 3D ECM scaffolds (type ECM-D) for HSC culture. Immunofluorescent staining of α-smooth muscle actin (αSMA) reveals activation of HSCs in standard 2D culture (top row) whereas HSCs in 3D matrix remain quiescent (bottom row). **C** Immunofluorescent staining of vimentin and αSMA shows a quiescent (αSMA negative) HSC phenotype in the untreated Quad Liver-Chip. **D** Representative images of AdipoRed staining showing lipid droplet accumulation (left) and quantification of lipid droplet count per cell and size (right) in the Quad Liver-Chip in response to either ethanol alone or ethanol + LPS. **E** Fold-change of lipid droplet count per cell and size (compared to median value of untreated chips) in Quad Liver-Chips and Tri Liver-Chips. Data are from one experiment with n=2-3 chips per condition. Data represent average ± (min and max). \**p*< 0.05; \*\**p*< 0.01; \*\*\**p*< 0.001; \*\*\*\**p*< 0.0001 versus control (ANOVA and Sidak’s test).

## Notes

### Competing Interest Statement

JCN, DBP, DVM, AS, GH, KK are current or former employees and may hold equity of Emulate, Inc.

### Summary of Updates

Clarified sentence in abstract Fixed typo in author list Updated affiliation

